# Effect of using a physical assistance device for movements involving trunk bending

**DOI:** 10.1101/2021.02.08.429597

**Authors:** Z. Jelti, K. Lebel, S. Bastide, P. Le Borgne, P. Slangen, N. Vignais

## Abstract

Exoskeletons are a solution to physically relieve workers while allowing them to control the execution of their tasks and assist them *(Baltrusch et al.2018)*.

The study investigated in the search for elements able to quantify the action of a physical assistance device (PAD) on the human body for movements responsible for pathologies recognized as occupational diseases.

The evaluation of a PAD allows to determine in which framework the exoskeleton can be useful for the realization of the movement. It is specified that a relevant way to insert exoskeletons in a company by always implementing a study or analysis beforehand to ensure its good integration.

The objective of the study is to perform several tasks with and without the Posture Harness (HAPO) in order to analyze the kinematics of the movements and the electrical activity of erector muscles of the spine involved in carrying a load at work to understand its effects on the human body.

## INTRODUCTION

Work-related musculoskeletal disorders (MSDs) are the leading cause of occupational diseases. MSDs have multiple causes. Among them, occupational activity frequently plays a role in their onset, maintenance or aggravation. MSDs result from an imbalance between the body’s physical capacities and the stresses and strains to which the body is exposed. MSDs usually develop gradually after a long period of intensive use of the affected parts of the body.

Absences from work due to MSDs can affect the performance of companies in which material handling plays an important role.

In the professional world, more precisely in logistics and industry, the issue of exoskeletons is their acceptance by the user and their adequacy with the work environment. This is the reason why companies are partnering with PAD designers in order to obtain devices that are faithful to the different workstations. Before deploying a device, it is important to make sure that it works properly and is reliable. To address these issues, a generalized device for MSD prevention should be designed, including lifting assistance able to relieve the lumbar area. PADs are designed to respect the biomechanics of human movement, however there are still important scientific locks. Understanding the control of human movement during interaction with an exoskeleton is a crucial step.

The Institute National for Research and Security (INRS, France) follows this approach in its study on exoskeletons at work *(Theurel and Schwartz, 2020)*. They compare different physical assistance devices during several lifting tasks. They underline the importance of clearly identifying the task that one wishes to assist in order to choose a relevant exoskeleton. Thus, in order to obtain positive results, the chosen physical assistance device must be specific to the task.

Based on the knowledge available in the literature, it seems that exoskeletons are effective for specific tasks such as arm elevation, sagittal trunk extension. In their work, the INRS shows a 10 to 80% decrease in the activity of the muscles targeted by the assistance. Such discrepancies can be explained by the fact that exoskeletons are not always well adapted to the situation in which they are used. In addition, the morphological and motor variability of users is not always taken into account in the design of such devices *(Theurel and* Schwartz, *2020)*.

The limitations outlined above often came from a lack of understanding of human-exoskeletal interactions. For this reason, it is essential to choose the device according to the desired assistance and the specificities of the task. For this, a first necessary step is to analyze precisely the action of the device on the user during a situation near to a real situation.

## PROTOCOL

A preliminary study at the laboratory of the Institute Mines – Telecom of Alès using motion capture (Qualisys cameras, miqus 3) has enabled us to identify the parameters necessary for our study. Objective measurements on the stability of the standing posture, lumbar spinal deformation and the muscular activity of the erector muscles of the user’s spine were therefore analyzed.

### Participants

Twelve healthy participants (7 men and 5 women), with an average age of 23.14 years (s=±2.51), an average weight of 73.71 kg (s=± 17.52) and an average height of 1.73 m (s=± 0.12 m), carried out in an experiment in the COGITOBIO movement analysis laboratory. Volunteer participants were required to complete an informed consent form prior to the experimentation.

The research protocol was checked and validated by the Research Ethics Committee (REC) POLETHIS of the University Paris Saclay (France). Participants have no documented background of locomotion and balance disorders or medical conditions that could affect postural control over the past two years.

### Materials

The posture harness evaluated in this study is the HAPO, developed, designed by Ergosanté (France) and marketed in March 2019. The HAPO is a lightweight (1.2 kg) physical assistance device that provides torque support for the user’s back, transferring the stresses to the sleeves of the thighs.

Objective measurements were carried out using ground reaction force (AMTI force platform 60cm, frequency 1000Hz), surface electromyography (EMG, Waves Plus Cometa, frequency 1000Hz) on the right longissimus muscle and the biceps femoris muscle. Inertial Measurements Units (Cogitobio, frequency 1000Hz) were placed at T1, T11 and L5 to measure lumbar lordosis. Subjective assessments of the perception and acceptance of the device were also carried out taking into account the INRS recommendations *(Wioland et al, 2019)*.

### Methods

The participants performed a static task of bending the trunk at 45 degrees with the arms extended forward for 2 minutes and several repetitions of a lifting task of 7.5 kg under two conditions “Without HAPO” and “With HAPO engaged” in the most natural way possible.

After completing the load carrying task under the “HAPO engaged” condition, the participants completed a questionnaire to assess the acceptance of the PAD.

### Data analysis

The data processing was carried out using the Python 3 programming language. After checking the normality of our samples (Shapiro-Wilk test), statistical tests were performed for each of the parameters studied (Wilcoxon test). For all the statistical tests, the significance threshold was set at α = 0.05.

## RESULTS ET DISCUSSION

### Feelings questionnaire

The questionnaire was completed at the end of each “HAPO engaged” condition. The satisfaction rate of the various participants is shown in figure 5. The criteria for judging the use of the HAPO are “Ease to use”, “Ease to implement”, “Effort reduction”, “Good comfort”, “Effectiveness of the system”.

**Figure 1:**
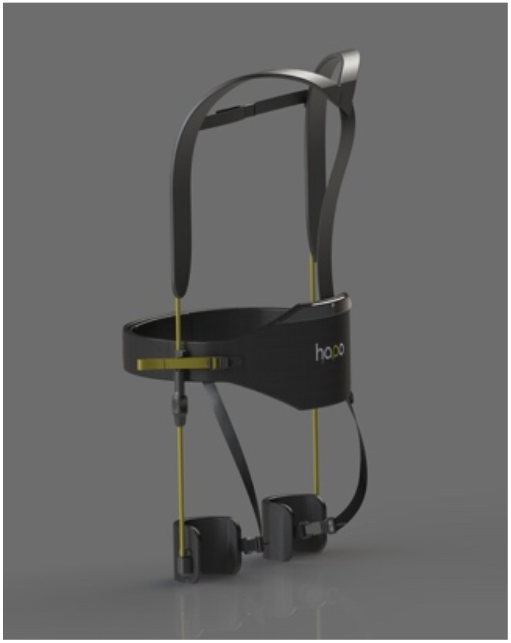
The Physical Assistance Device HAPO, posture harness.

**Figure 2:**
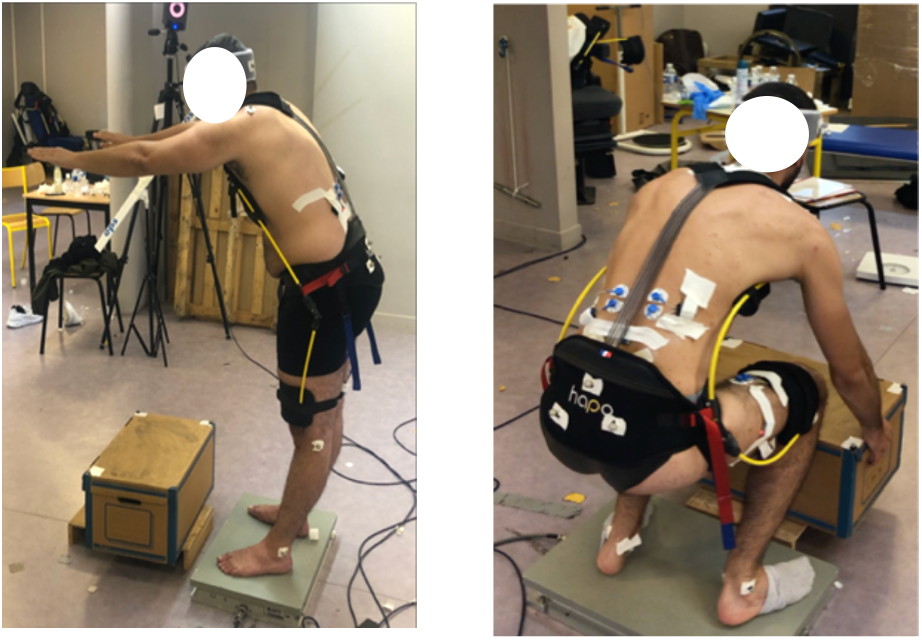
Photography of the experiment, on the right a participant carrying out the load carrying task, on the left a participant carrying out the static task.

**Figure 3:**
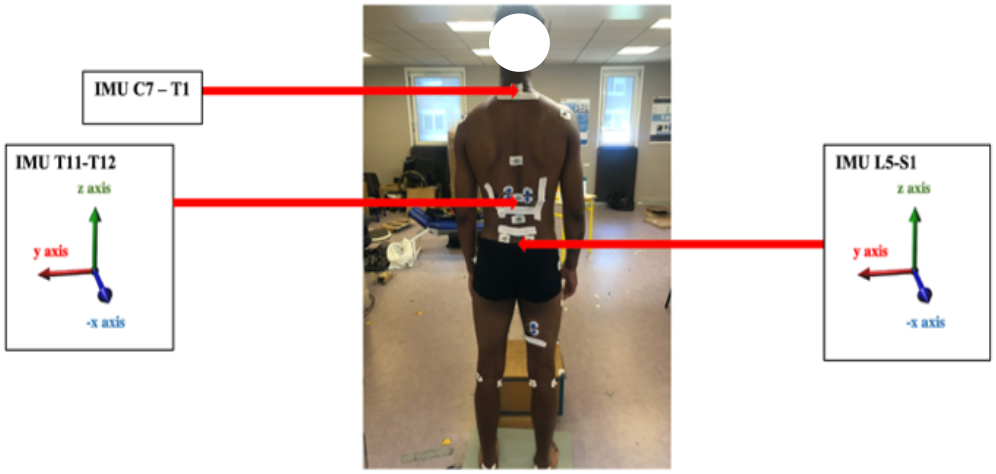
Arrangements of the inertial measurement units on the T1, T11 and L5 vertebrae.

**Figure 4:**
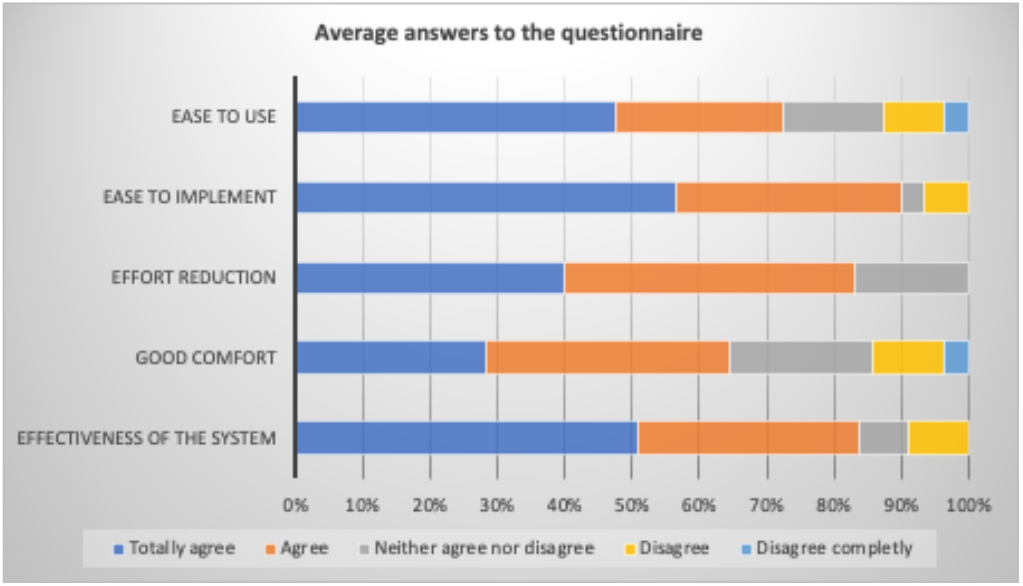
Mean answers to the feelings and comfort questionnaire

**Figure 5:**
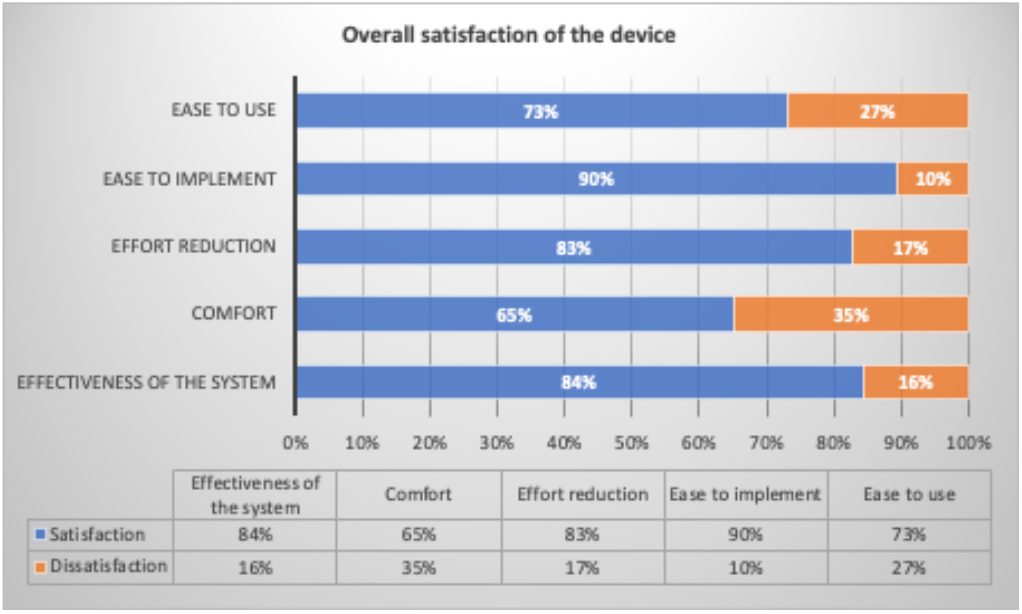
Mean answers to the Feelings and Comfort questionnaire represented as satisfaction.

**Figure 6:**
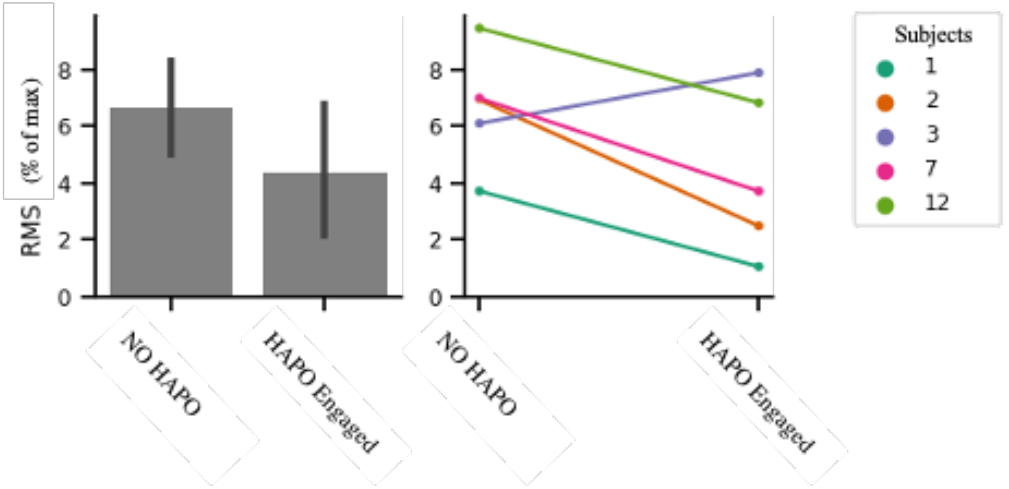
Root Mean Square mean of muscle activity of the right longissimus muscle.

**Table 1:** Descriptive statistics of the area of the 95% confidence ellipse.

|  | Mean (mm<sup>2</sup>) | Standard deviation |
| --- | --- | --- |
| Area of CoP No HAPO | 1033.15 | 632.37 |
| Area of CoP HAPO engaged | 857.95 | 1003.97 |

**Figure 7:**
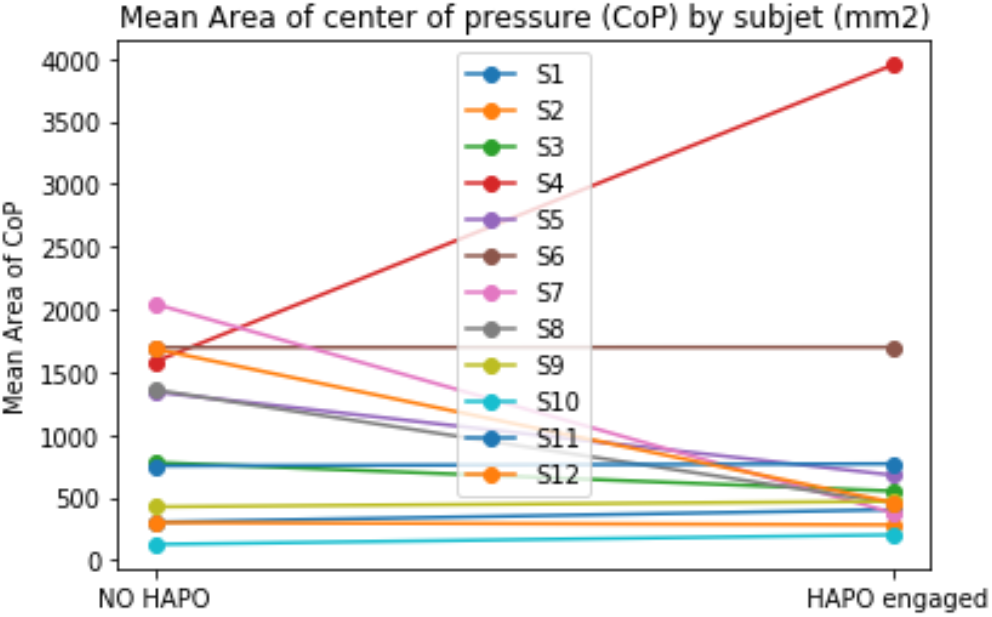
Mean area of the 95% confidence ellipse “With HAPO engaged” and “Without HAPO”.

**Figure 8:**
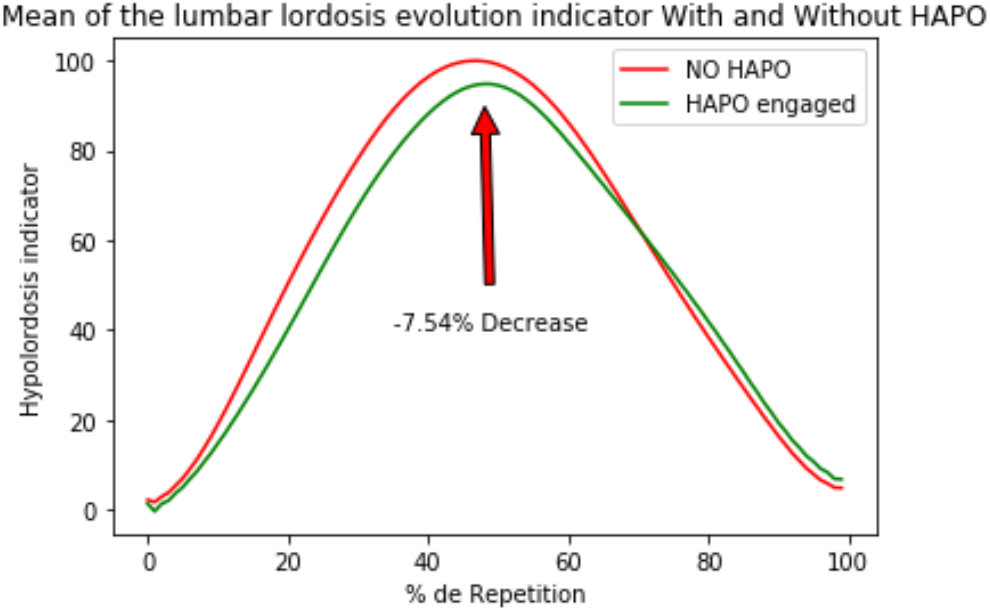
Mean curve of the lumbar hypolordosis indicator with HAPO engaged and without HAPO

For a lifting task, the HAPO satisfaction of the participants during the experiment reached a rate of 79 ± 9.92%. This result shows that the participants had a good feeling when using the HAPO.

Moreover, its effectiveness was felt from the first trunk extension movement.

### Electromyography (EMG)

The literature tells us that PADs reduce the activity of the spinal muscles. In this study, the analysis focuses on the longissimus muscle. EMG measurements were analyzed on five participants. The aim was to quantify the overall muscle contraction of each individual.

Root Mean Square – Right longissimus muscle

Trends can be observed: the use of HAPO makes it possible, as expected thanks to the work of *Baltrusch et al (2018)* and *Theurel and Desbrosses (2019)*, to reduce the muscular activity of the erectors of the rachis (here the longissimus muscle) by an average of 33.8%. A decrease in the activity of the long biceps femoral muscle was also observed. This relieves hip and knee extension during a lifting task.

Both the activation peak during the movement and the total amount of activation decreases.

This could have beneficial consequences on muscle fatigue and the risk of long-term musculoskeletal disorders.

### Force plate

The force platform was used to obtain the displacement of the center of pressure during the different tasks carried out. This parameter allows us to quantify the postural balance between each condition.

We can see that the postural balance goes towards a decrease with the condition “With HAPO engaged”. Moreover, in the condition with harness, we observe that there is less variation. The majority of subjects seem to have better balance control with the PAD. However, in our study, this balance control varied from one subject to another, which does not allow us to statistically validate that there is a significant difference between the two conditions (p > 0.05). It can therefore be concluded that there is no significant change from one condition to another although there is a decrease for some of the participants.

### « Spinometer »: Inertial Measurements Units

The analysis of inertial units was carried out on nine participants.

Concerning the deformation of the spinal column in lumbar lordosis, it was found that there is a significant change from one condition to another (p < 0.05). The engaged HAPO reduces the change in the curvature of the lumbar spine for tasks involving trunk bending.

A phenomenon of modification of the lumbar lordosis can be observed. This is an average reduction of 7.54 ± 4.55 % in the lumbar concavity hypolordosis for a lifting task. This can be as much as a 16% reduction in some subjects. Lifting with HAPO appears not to disturb the articular blocks or chains of the spine, which therefore does not alter the kinematics of movement. A much more sustained study using clinical radiology equipment would validate precisely these analyses.

## CONCLUSION

Through this experiment, we observed that the use of the HAPO, posture harness, for tasks requiring bending trunk, reduced the muscular solicitation of the erector muscles of the spine while reducing hypolordosis.

These results show that the exoskeleton could make it possible to preserve the integrity of the physiological curvature of the lumbar lordosis of the spine. The shear stresses of the intervertebral discs would therefore be limited. Moreover, these improvements occur without any adverse effect on postural balance.

Overall, the HAPO exoskeleton has a beneficial effect on muscle activity and spinal curvature. Despite non-significant results, results on a larger number of subjects could demonstrate the interest of HAPO for postural control.

This experiment allows us to better understand the preventive effects of HAPO on users.

This employed approach is promising for evaluating exoskeletons and attesting to their effectiveness.

## ACKNOWLEDGEMENTS

This study was made possible thanks to a collaboration between the CIAMS laboratory, COGITOBIO laboratory, CERIS laboratory and the company ERGOSANTÉ.

